# Modeling the language of life – Deep Learning Protein Sequences

**DOI:** 10.1101/614313

**Authors:** Michael Heinzinger, Ahmed Elnaggar, Yu Wang, Christian Dallago, Dmitrii Nechaev, Florian Matthes, Burkhard Rost

## Abstract

**Background:** One common task in Computational Biology is the prediction of aspects of protein function and structure from their amino acid sequence. For 26 years, most state-of-the-art approaches toward this end have been marrying machine learning and evolutionary information. The retrieval of related proteins from ever growing sequence databases is becoming so time-consuming that the analysis of entire proteomes becomes challenging. On top, evolutionary information is less powerful for small families, e.g. for proteins from the *Dark Proteome*.

**Results:** We introduce a novel way to represent protein sequences as continuous vectors (*embeddings*) by using the deep bi-directional model ELMo taken from natural language processing (NLP). The model has effectively captured the biophysical properties of protein sequences from unlabeled big data (UniRef50). After training, this knowledge is transferred to single protein sequences by predicting relevant sequence features. We refer to these new embeddings as *SeqVec* (*Seq*uence-to-*Vec*tor) and demonstrate their effectiveness by training simple convolutional neural networks on existing data sets for two completely different prediction tasks. At the per-residue level, we significantly improved secondary structure (for NetSurfP-2.0 data set: Q3=79%±1, Q8=68%±1) and disorder predictions (MCC=0.59±0.03) over methods not using evolutionary information. At the per-protein level, we predicted subcellular localization in ten classes (for DeepLoc data set: Q10=68%±1) and distinguished membrane-bound from water-soluble proteins (Q2= 87%±1). All results built upon the embeddings gained from the new tool *SeqVec* neither explicitly nor implicitly using evolutionary information. Nevertheless, it improved over some methods using such information. Where the lightning-fast *HHblits* needed on average about two minutes to generate the evolutionary information for a target protein, *SeqVec* created the vector representation on average in 0.03 seconds.

**Conclusion:** We have shown that transfer learning can be used to capture biochemical or biophysical properties of protein sequences from large unlabeled sequence databases. The effectiveness of the proposed approach was showcased for different prediction tasks using only single protein sequences. *SeqVec* embeddings enable predictions that outperform even some methods using evolutionary information. Thus, they prove to condense the underlying principles of protein sequences. This might be the first step towards competitive predictions based only on single protein sequences.

**Availability:** SeqVec: https://github.com/mheinzinger/SeqVec Prediction server: https://embed.protein.properties

## Background

Over two decades ago, the combination of evolutionary information (from Multiple Sequence Alignments – MSA) and machine learning (standard feed-forward artificial neural networks – ANN) completely changed protein secondary structure prediction [1-3]. The concept was quickly taken up [4-8], and it was shown how the improvement in prediction increased with even larger families including more diverse evolutionary information [9, 10]. The idea was applied to other tasks, including the prediction of transmembrane regions [11-13], solvent accessibility [14], residue flexibility (B-values) [15, 16], inter-residue contacts [17] and protein disorder [15, 18-20]. Later, methods predicting aspects of protein function improved through the combination of evolutionary information and machine learning, including predictions of sub-cellular localization (aka cellular location [21, 22]), protein interaction sites [23-25], and the effects of sequence variation upon function [26, 27]. Arguably, the most important breakthrough for protein structure prediction over the decade was a more efficient way of using evolutionary couplings [28-31].

Although evolutionary information has become increasingly crucial, it is also becoming increasingly costly. Firstly, bio-sequence databases grow faster than computers making it expensive to find and align related proteins. For instance, the number of UniProt entries is now more than doubling every two years [32]. Consequently, methods as fast as PSI-BLAST [33] have to be replaced by faster solutions such as HHblits [34]. Even its latest version HHblits3 [35] still needs several minutes to search UniRef50 (subset of UniProt; ∼20% of UniProt release 2019_02). The next step up in speed such as MMSeqs2 [36] appear to cope with the challenge at the expense of increasing hardware requirements while databases keep growing. However, even these solutions might eventually lose the battle against the speedup of sequencing. Analyzing data sets involving millions of proteins, i.e. samples of the human gut microbiota or metagenomic samples, will be one of the major challenges for computational biology. Secondly, evolutionary information is missing for some proteins, e.g. for proteins with substantial intrinsically disordered regions [15, 37, 38], or the entire *Dark Proteome* [39] full of proteins that are less-well studied but important for function [40].

Here, we propose a novel embedding of protein sequences that replaces the explicit search for evolutionary related proteins by an implicit transfer of biophysical information derived from large, unlabeled sequence data (here UniRef50). Towards this end, we have adopted a method that has been revolutionizing Natural Language Processing (NLP), namely the bidirectional language model ELMo (Embeddings from Language Models) [41]. In NLP, ELMo is trained on unlabeled text-corpora such as Wikipedia to predict the most probable next word in a sentence, given all previous words in this sentence. By learning a probability distribution for sentences, these models develop autonomously a notion for syntax and semantics of language. The trained vector representations (embeddings) are contextualized, i.e. embeddings of a given word depend on its context. This has the advantage that two identical words can have different embeddings, depending on the words surrounding them.

We hypothesized that the ELMo concept could be applied to learn aspects of what makes up the language of life distilled in protein sequences. Three main challenges arose. (1) Proteins range from about 30 to 33,000 residues, a much larger range than for the average English sentence extending over 15-30 words [42], and even more extreme than notable literary exceptions such as James Joyce’s Ulysses (1922) with almost 4,000 words in a sentence. Longer proteins require more GPU memory and the underlying models (so called Long Short-Term Memory networks (LSTMs) [43]) have only a limited capability to remember long-range dependencies. (2) Proteins mostly use 20 standard amino acids, 100,000 times less than in the English language. Smaller vocabularies might be problematic if protein sequences encode a similar complexity as sentences. (3) We found UniRef50 to contain almost ten times more tokens (9.5 billion amino acids) than the largest existing NLP corpus (1 billion words). Simply put: Wikipedia is roughly ten times larger than Webster’s Third New International Dictionary and the entire UniProt is over ten times larger than Wikipedia. As a result, larger models might be required to absorb the information provided.

We trained the bi-directional language model ELMo on UniRef50. Then we assessed the predictive power of the embeddings by application to tasks on two levels: per-residue (word-level) and per-protein (sentence-level). For the per-residue prediction task, we predicted secondary structure in three (helix, strand, other) and eight states (all DSSP [44]), as well as long intrinsic disorder in two states. For the per-protein prediction task, we implemented the predictions of protein subcellular localization in ten classes and a binary classification into membrane-bound and water-soluble proteins. We used publicly available data sets from two recent methods that achieved break-through performance through Deep Learning, namely NetSurfP-2.0 (secondary structure [45]) and DeepLoc (localization [46]). We compared the performance of the *SeqVec* embeddings to state-of-the-art methods, and also to a popular embedding tool for protein sequences, namely *ProtVec* [47]. ProtVec assumes that every token or word consists of three consecutive residues (amino acid 3-mers). During training, each protein sequence in *SwissProt* [48] is split into overlapping 3-mers and another tool from NLP, namely the *word2vec* (skip-gram) model [49] is used to predict adjacent words, given the word at the center. After training, protein sequences can be split into overlapping 3-mers which are mapped onto a 100-dimensional latent space. While this approach can capture local information, it loses information on sequence ordering and the resulting embeddings are insensitive to their context (non-contextualized), i.e. the same word results in the same embedding regardless of the specific context.

## Results

### *SeqVec* embeddings compress UniProt50

*SeqVec*, our ELMo-like implementation was trained for three weeks on 5 Nvidia Titan GPUs with 12 GB memory each. The model was trained until its *perplexity* (uncertainty when predicting the next token) converged at around 10.5 (Fig. SOM_1). Training and testing were not split due to technical limitations (incl. shortage of CPU/GPU). ELMo was designed to reduce the risk of overfitting by sharing weights between forward and backward LSTMs and by using dropout. The model had about 93M (mega/million) free parameters which was about 100-times smaller than the 9.6G (giga/billion) tokens to predict. Similar approaches have shown that even todays largest models (750M free parameters), are not able to overfit on a large corpus (250M protein sequences) [50].

### Per-residue performance high, not highest

NetSurfP-2.0 uses HHblits profiles along with advanced combinations of Deep Learning architectures [45]. NetSurfP-2.0 could be today’s best method for protein secondary structure prediction, reaching a three-state per-residue accuracy Q3 of 82-85% (lower value: small very non-redundant CASP12 set, upper value: larger slightly more redundant TS115 and CB513 sets; Table 1, Fig. 1; note that several contenders such as *Spider3* and *RaptorX* appear to differ by fewer than three standard errors). All six methods developed by us (DeepProf, DeepSeqVec, DeepProf+SeqVec, DeepProtVec, DeepOneHot, DeepBLOSUM65) fell short of reaching this mark (Fig. 1A, Table 1). When comparing methods that use only single protein sequences as input (DeepSeqVec, DeepProtVec, DeepOneHot, DeepBLOSUM65; all white in Table 1), the proposed *SeqVec* outperformed others by 5-10 (Q3), 5-13 (Q8) and 0.07-0.12 (MCC) percentage points. The evolutionary information (DeepProf with HHblits profiles) remained about 4-6 percentage points below NetSurfP-2.0 (Q3=76-81%, Fig. 1, Table 1). Depending on the test set, using *SeqVec* embeddings instead of evolutionary information (DeepSeqVec: Fig. 1A, Table 1) remained 2-3 percentage points below that mark (Q3=73-79%, Fig. 1A, Table 1). Using both evolutionary information and *SeqVec* embeddings (DeepProf+SeqVec) improved over both, but still did not reach the top (Q3=77-82%). In fact, the embedding alone (DeepSeqVec) did not surpass any of the existing methods using evolutionary information (Fig. 1A).

**Table 1:** Per-residue predictions: secondary structure and disorder ^◊^.

| Data | Prediction task | Secondary structure |  | Disorder |  |
| --- | --- | --- | --- | --- | --- |
|  | Method | Q3 (%) | Q8 (%) | MCC | FPR |
| CASP12 | NetSurfP-2.0 (hhblits)* | **82.4** | **71.1** | 0.604 | **0.011** |
|  | NetSurfP-1.0* | 70.9 | - | - | - |
|  | Spider3* | 79.1 | - | 0.582 | 0.026 |
|  | RaptorX* | 78.6 | 66.1 | **0.621** | 0.045 |
|  | Jpred4* | 76.0 | - | - | - |
|  | DeepSeqVec | 73.1 ± 1.3 | 61.2 ± 1.6 | 0.575 ± 0.075 | 0.026 ± 0.008 |
|  | DeepProf | 76.4 ± 2.0 | 62.7 ± 2.2 | 0.506 ± 0.057 | 0.022 ± 0.009 |
|  | DeepProf + SeqVec | 76.5 ± 1.5 | 64.1 ± 1.5 | 0.556 ± 0.080 | 0.022 ± 0.008 |
|  | DeepProtVec | 62.8 ± 1.7 | 50.5 ± 2.4 | 0.505 ± 0.064 | 0.016 ± 0.006 |
|  | DeepOneHot | 67.1 ± 1.6 | 54.2 ± 2.1 | 0.461 ± 0.064 | 0.012 ± 0.005 |
|  | DeepBLOSUM65 | 67.0 ± 1.6 | 54.5 ± 2.0 | 0.465 ± 0.065 | 0.012 ± 0.005 |
| TS115 | NetSurfP-2.0 (hhblits)* | **85.3** | **74.4** | **0.663** | **0.006** |
|  | NetSurfP-1.0* | 77.9 | - | - | - |
|  | Spider3* | 83.9 | - | 0.575 | 0.008 |
|  | RaptorX* | 82.2 | 71.6 | 0.567 | 0.027 |
|  | Jpred4* | 76.7 | - | - | - |
|  | DeepSeqVec | 79.1 ± 0.8 | 67.6 ± 1.0 | 0.591 ± 0.028 | 0.012 ± 0.001 |
|  | DeepProf | 81.1 ± 0.6 | 68.3 ± 0.9 | 0.516 ± 0.028 | 0.012 ± 0.002 |
|  | DeepProf + SeqVec | 82.4 ± 0.7 | 70.3 ± 1.0 | 0.585 ± 0.029 | 0.013 ± 0.003 |
|  | DeepProtVec | 66.0 ± 1.0 | 54.4 ± 1.3 | 0.470 ± 0.028 | 0.011 ± 0.002 |
|  | DeepOneHot | 70.1 ± 0.8 | 58.5 ± 1.1 | 0.476 ± 0.028 | 0.008 ± 0.001 |
|  | Deep BLOSUM65 | 70.3 ± 0.8 | 58.1 ± 1.1 | 0.488 ± 0.029 | 0.007 ± 0.001 |
| CB513 | NetSurfP-2.0 (hhblits)* | **85.3** | **72.0** | - | - |
|  | NetSurfP-1.0* | 78.8 | - | - | - |
|  | Spider3* | 84.5 | - | - | - |
|  | RaptorX* | 82.7 | 70.6 | - | - |
|  | Jpred4* | 77.9 | - | - | - |
|  | DeepSeqVec | 76.9 ± 0.5 | 62.5 ± 0.6 | - | - |
|  | DeepProf | 80.2 ± 0.4 | 64.9 ± 0.5 | - | - |
|  | DeepProf + SeqVec | 80.7 ± 0.5 | 66.0 ± 0.5 | - | - |
|  | DeepProtVec | 63.5 ± 0.4 | 48.9 ± 0.5 | - | - |
|  | DeepOneHot | 67.5 ± 0.4 | 52.9 ± 0.5 | - | - |
|  | DeepBLOSUM65 | 67.4 ± 0.4 | 53.0 ± 0.5 | - | - |

\* Performance comparison for 3- and 8-class secondary structure prediction as well as disorder prediction for the CASP12, TS115 and CB513 data sets. Accuracies (Q3, Q10) are given in percentage. Results marked with \* are taken from NetSurfP-2.0 [45]; the authors did not provide standard errors. DeepSeqVec, DeepProtVec, DeepOneHot and DeepBLOSUM65 use only information from single protein sequences. All methods using evolutionary information are marked by gray boxes, these performed best (best in each set marked by bold numbers) for all tasks and measures.

**Figure 1:**
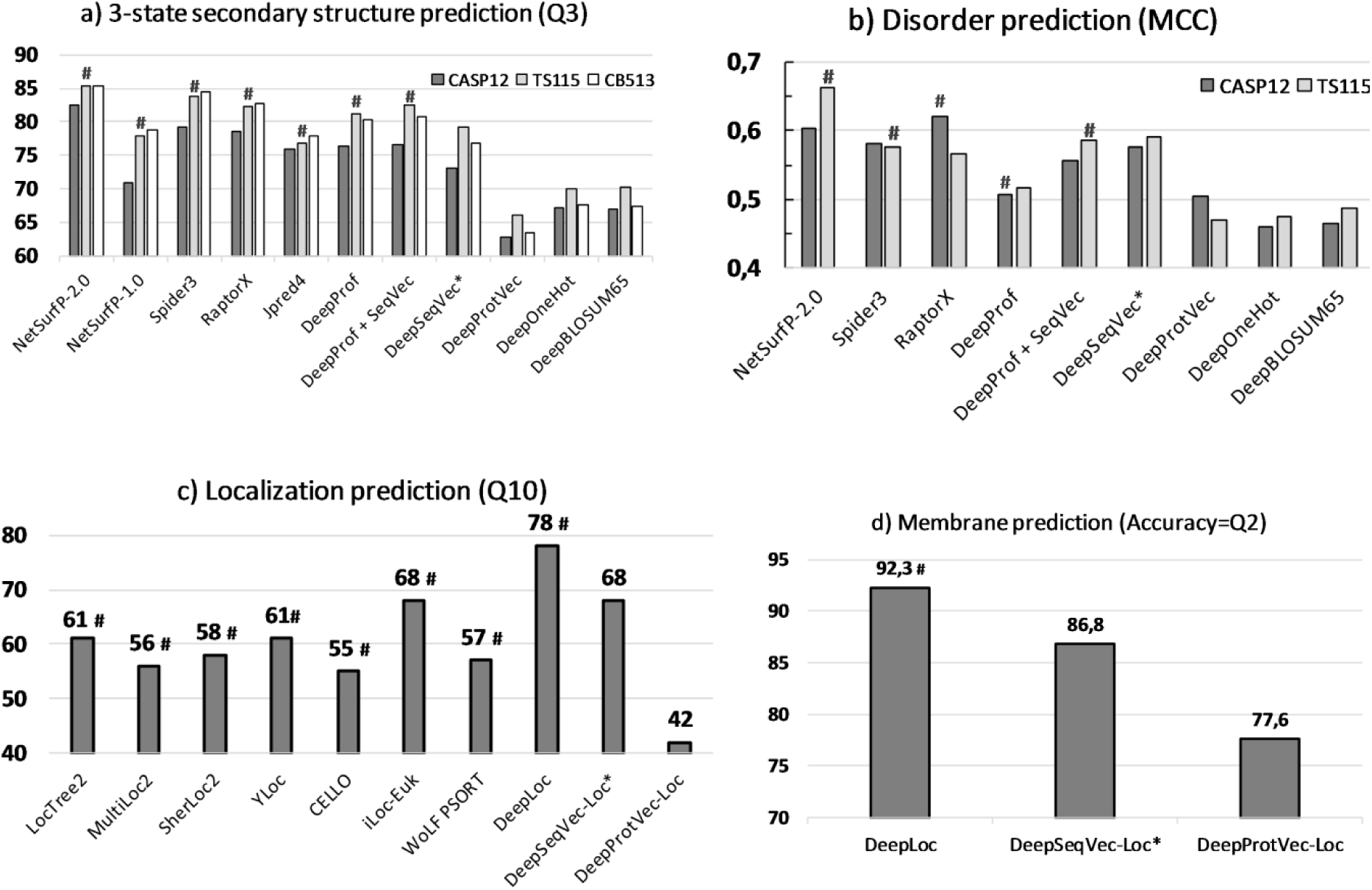
Performance comparisons. The predictive power of the ELMo-like SeqVec embeddings was assessed for per-residue (upper row) and per-protein (lower row). Methods using evolutionary information (mostly in the form of alignments) are highlighted by a ‘#’ above the bar(s) of the methods. Approaches using only the proposed SeqVec embeddings are highlighted by a ‘*’. <u>Panel A</u> compared three-state secondary structure prediction of the proposed SeqVec to other embeddings based on single protein sequences. <u>Panel B</u> compared predictions of intrinsically disordered residues. <u>Panel C</u> compared per-protein predictions for subcellular localization between top methods (numbers taken from DeepLoc [46] and embeddings based on single sequences (ProtVec [47] and our SeqVec). <u>Panel D</u>: the same data set was used to assess the predictive power of SeqVec for the classification of a protein into membrane-bound and water-soluble.

For the prediction of intrinsic disorder, we observed the same: NetSurfP-2.0 performed best; our implementation of evolutionary information (DeepProf) performed worse (Fig. 1B, Table 1). However, for this task the embeddings alone (DeepSeqVec) performed relatively well, exceeding our in-house implementation of a model using evolutionary information (DeepSeqVec MCC=0.575-0.591 vs. DeepProf MCC=0.506-0.516, Table 1). The combination of evolutionary information and embeddings (DeepProf+SeqVec) improved over using evolutionary information alone but did not improve over the *SeqVec* embeddings for disorder. Compared to other methods, the embeddings alone reached similar values (Fig. 1B).

### Per-protein performance close to best

For the task of predicting subcellular localization in ten classes, DeepLoc [46] appears to be today’s best tool with Q10=78% (Fig. 1C, Table 2). Our simple embeddings model DeepSeqVec-Loc reached second best performance together with iLoc-Euk [51] with Q10=68% (Fig. 1C, Table 2). For this application the SeqVec embeddings outperformed several other methods that use evolutionary information by up to Q10=13%.

**Table 2:** Per-protein predictions: localization and membrane/globular ^◊^.

| Method | Localization |  | Membrane/globular |  |
| --- | --- | --- | --- | --- |
|  | Q10 (%) | Gorodkin (MCC) | Q2 | MCC |
| LocTree2* | 61 | 0.53 |  |  |
| MultiLoc2* | 56 | 0.49 |  |  |
| CELLO* | 55 | 0.45 |  |  |
| WoLF PSORT* | 57 | 0.48 |  |  |
| YLoc* | 61 | 0.53 |  |  |
| SherLoc2* | 58 | 0.51 |  |  |
| iLoc-Euk* | 68 | 0.64 |  |  |
| DeepLoc* | **78** | **0.73** | **92.3** | **0.844** |
| DeepSeqVec-Loc | 68 ± 1 | 0.61 ± 0.01 | 86.8 ± 1.0 | 0.725 ± 0.021 |
| DeepProtVec-Loc | 42 ± 1 | 0.19 ± 0.01 | 77.6 ± 1.3 | 0.531 ± 0.026 |

<sup>◇</sup> Performance for per-protein prediction of subcellular localization and the classification of proteins into membrane-bound and water-soluble. Results marked with \* were taken from DeepLoc [46]; the authors did not provide standard errors. The results reported for SeqVec and ProtVec were based on single protein sequences, i.e. methods NOT using evolutionary information (neither during training nor testing). All methods using evolutionary information are marked by gray boxes, these performed best (best in each set marked by bold numbers) for all tasks and measures.

Performance for the classification into membrane-bound and water-soluble proteins followed a similar trend (Fig. 1D, Table 2): while DeepLoc still performed best (Q2=92.3, MCC=0.844), DeepSeqVec-Loc reached just a few percentage points lower (Q2=86.8±1.0, MCC=0.725±0.021; full confusion matrix Figure SOM_2). In contrast to this, ProtVec, another method using only single sequences, performed substantially worse (Q2=77.6±1.3, MCC=0.531±0.026).

### Visualizing results

Lack of insight often triggers the misunderstanding that machine learning methods are black box solutions barring understanding. In order to interpret the *SeqVec* embeddings, we have projected the protein-embeddings of the per-protein prediction data upon two dimensions using t-SNE [52]. We performed this analysis once for the raw embeddings (SeqVec, Fig. 2 upper row) and once for the hidden layer representation of the per-protein network (DeepSeqVec-Loc) after training (Fig. 2 lower row). All t-SNE representations in Fig. 2 were created using 3,000 iterations and the cosine distance as metric. The two analyses differed only in that the perplexity was set to 20 for one (*SeqVec*) and 15 for the other (DeepSeqVec-Loc). The t-SNE representations were colored either according to their localization within the cell (left column of Fig. 2) or according to whether they are membrane-bound or water-soluble (right column).

**Figure 2:**
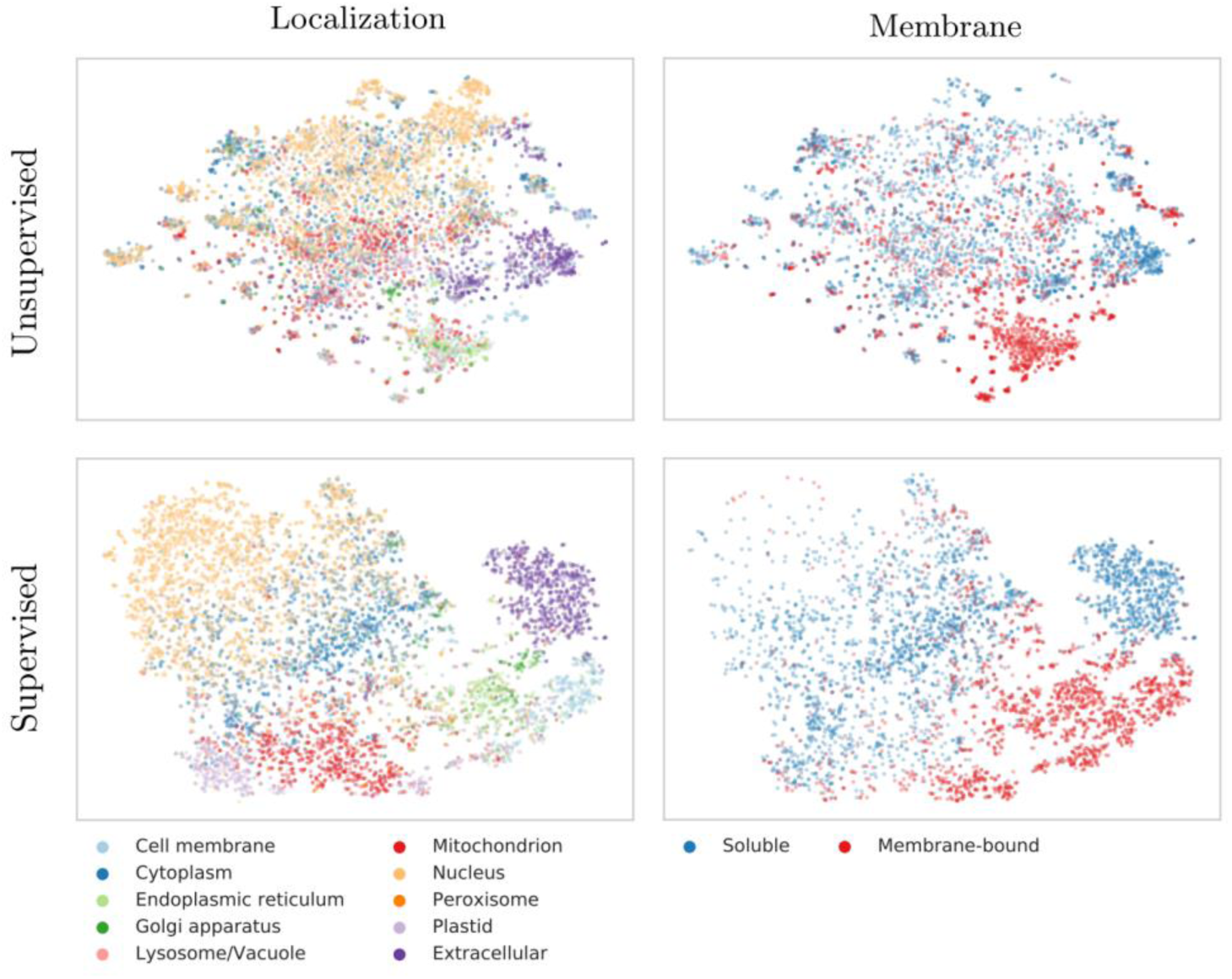
t-SNE representations of SeqVec. t-SNE projections from embedded space onto a 2D representation; upper row: unsupervised 1024-dimensional ELMo embeddings, averaged over all residues in a protein; lower row: supervised 32 dimensional ELMo embeddings, reduced via per-protein machine learning predictions. The redundancy reduced DeepLoc data set was used for this figure. Proteins were colored according to their localization (left column) or whether they are membrane-bound or water-soluble (right column). Not all proteins in the data set have annotations for the classification into membrane-bound/water-soluble and have thus been excluded in the panels on the right. The plots in the upper row suggest that the SeqVec embeddings already capture various aspects of proteins prior even to supervised learning. After supervised training (lower row), this information is transferred to, and further distilled by networks with simple architectures to distinguish aspects of protein function (sometimes drastically, as in the case of membrane boundness, as suggested by the coloured clusters in the bottom right panel). After training machine learning devices (lower row) on the supervised tasks of localization (left) and membrane boundness (right) prediction, the power of SeqVeq embeddings to distinguish aspects of function and structure become even more pronounced, sometimes drastically so, as suggested by the almost fully separable clusters in the lower right panel.

Despite never provided during training, the raw embeddings appeared to capture some signal for classifying proteins by localization (Fig. 2, upper row, left column). The most consistent signal was visible for extra-cellular proteins. Proteins attached to the cell membrane or located in the endoplasmic reticulum also formed well-defined clusters. In contrast, the raw embeddings neither captured an ambiguous signal for nuclear nor for mitochondrial proteins. Through training, the network improved the signal to reliably classify mitochondrial and plastid proteins. However, proteins in the nucleus and cell membrane continued to be poorly distinguished via t-SNE.

Coloring the t-SNE representations for membrane-bound or water-soluble proteins (Fig. 2, right column), revealed that the raw embeddings already provided well-defined clusters although never trained on membrane prediction (Fig. 2, upper row). After training, the classification was even better (Fig. 2, lower row).

Analogously, we used t-SNE projections to analyze SeqVec embeddings on different levels of complexity inherent to proteins (Fig. 3), ranging from the building blocks (amino acids, Fig. 3a), to secondary structure defined protein classes (Fig. 3b), over functional features (Fig. 3c), and onto the macroscopic level of the kingdoms of life and viruses (Fig. 3d; classifications in panels 3b-3d based on SCOPe [53]). Similar to the results described in [50], our projection of the embedding space confirmed that the model successfully captured bio-chemical and bio-physical properties on the most fine-grained level, i.e. the 20 standard amino acids (Fig. 3a). For example, aromatic amino acids (W, F, Y) are well separated from aliphatic amino acids (A, I, L, M, V) and small amino acids (A, C, G, P, S, T) are well separated from large ones (F, H, R, W, Y). The projection of the letter indicating an unknown amino acid (X), clustered closest to the amino acids alanine (A) and glycine (G) (data not shown). The two amino acids with the smallest side chains might be least biased towards other biochemical features like charge and they are the 2^nd^ (A) and 4^th^ (G) most frequent amino acids in our training set (Table SOM_1). Rare (O, U) and ambiguous amino acids (Z, B) were removed from the projection as their clustering showed that the model could not learn a reasonable embedding from the very small number of samples.

**Figure 3:**
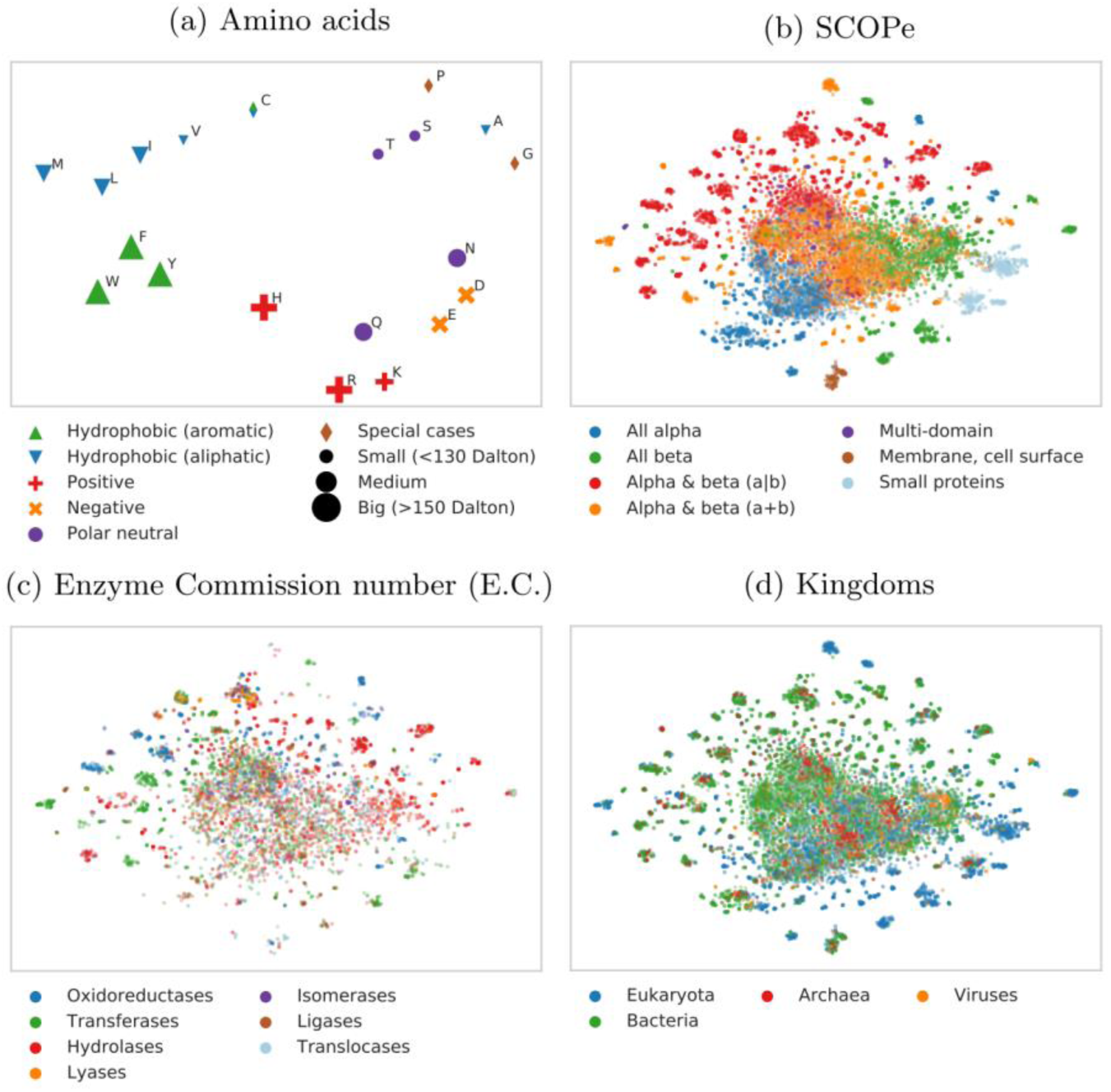
Modeling the language of life. 2D t-SNE projections of unsupervised SeqVec embeddings highlight different realities of proteins and their constituent parts, amino acids. Panels (b) to (d) are based on the same data set (Structural Classification of Proteins – extended (SCOPe) 2.07, redundancy reduced at 40%). For these plots, only subsets of SCOPe containing proteins with the annotation of interest (enzymatic activity (c) and kingdom (d)) may be displayed. Panel (a): the embedding space confirms: the 20 standard amino acids are clustered according to their biochemical and biophysical properties, i.e. hydrophobicity, charge or size. The unique role of Cysteine (C, mostly hydrophobic and polar) is conserved. Panel (b): SeqVec embeddings capture structural information as annotated in the main classes in SCOPe without ever having been explicitly trained on structural features. Panel (c): many small, local clusters share function as given by the main classes in the Enzyme Commission Number (E.C.). Panel (d): similarly, small, local clusters represent different kingdoms of life.

Crude structural classes as defined in SCOPe (Fig. 3b) were also captured by SeqVec embeddings. Although the embeddings were only trained to predict the next amino acid in a protein sequence, well separated clusters emerged from those embeddings in structure space. Especially, membrane proteins and small proteins formed distinct clusters (note: protein length did not affect the t-SNE grouping). Also, these results indicated that the embeddings captured complex relationships between proteins which are not directly observable from sequence similarity alone as SCOPe was redundancy reduced at 40% sequence identity. Therefore, the new embeddings could complement sequence-based structural classification as it was shown that the sequence similarity does not necessarily lead to structural similarity [54].

To further investigate the clusters which emerged from the SCOPe data set, we colored the same data set based on protein functions (Fig. 3c) and kingdoms (Fig. 3d). This analysis revealed that many of the small, distinct clusters emerged based on protein functions. For example, transferases and hydrolases form many small clusters. When increasing the level of abstraction by coloring the proteins according to their kingdoms, we observed certain clusters which are dominated by e.g. eukaryotes. Comparing the different views captured in panels 3b-3 revealed connections, e.g. that all-beta or small proteins dominate in eukaryotes (compare blue and orange islands in Fig. 3b with the same islands in Fig. 3d – colored blue to mark eukaryotes).

### CPU/GPU time used

Due to the sequential nature of LSTMs, the time required to embed a protein grew linearly with protein length. Depending on the available main memory or GPU memory, this process could be massively parallelized. To optimally use available memory, batches are typically based on tokens rather than on sentences. Here, we sorted proteins according to their length and created batches of ≤15K tokens that could still be handled by a single Nvidia GeForce GTX1080 with 8GB VRAM. On average the processing of a single protein took 0.027s when applying this batch-strategy to the NetSurfP-2.0 data set (average protein length: 256 residues, i.e. shorter than proteins for which 3D structure is not known). The batch with the shortest proteins (on average 38 residues, corresponding to 15% of the average protein length in the whole data set) required about one tenth (0.003s per protein, i.e. 11% of that for whole set). The batch containing the longest protein sequences in this data set (1578 residues on average, corresponding to 610% of average protein length in the whole data set), took about six times more (1.5s per protein, i.e. 556% of that for whole set). When creating SeqVec for the DeepLoc set (average length: 558 residues; as this set does not require a 3D structure, it provides a more realistic view on the distribution of protein lengths), the average processing time for a single protein was 0.08 with a minimum of 0.006 for the batch containing the shortest sequences (67 residues on average) and a maximum of 14.5s (9860 residues on average). Roughly, processing time was linear with respect to protein length. On a single Intel i7-6700 CPU with 64GB RAM, processing time increased by roughly 50% to 0.41s per protein, with a minimum and a maximum computation time of 0.06 and 15.3s, respectively. Compared to an average processing time of one hour for 1000 proteins when using evolutionary information directly [45], this implied an average speed up of 120-fold on a single GeForce GTX1080 and 9-fold on a single i7-6700 when predicting structural features; the inference time of DeepSeqVec for a single protein is on average 0.0028s.

## Discussion

### ELMo alone did not suffice for top performance

On the one hand, none of our implementations of ELMo reached anywhere near today’s best (NetSurfP-2.0 for secondary structure and protein disorder and DeepLoc for localization and membrane protein classification; Fig. 1, Table 1, Table 2). Clearly, “just” using ELMo did not suffice to crack the challenges. On the other hand, some of our solutions appeared surprisingly competitive given the simplicity of the architectures. In particular for the per-protein predictions, for which *SeqVec* clearly outperformed the previously popular ProtVec [47] approach and even commonly used expert solutions (Fig. 1, Table 2: no method tested other than the top-of-the-line DeepLoc reached higher numerical values). For that comparison, we used the same data sets but could not rigorously compare standard errors which were unavailable for other methods. Estimating standard errors for our methods suggested that the differences were statistically significant as the difference was more than 7 sigmas for all methods (except DeepLoc (Q10=78) and iLoc-Euk(Q10=68)). The results for localization prediction implied that frequently used methods using evolutionary information (all marked with stars in Table 2) did not clearly outperform our simple ELMo-based tool (DeepSeqVec-Loc in Table 2). This was very different for the per-residue prediction tasks: here almost all top methods using evolutionary information numerically outperformed the simple model built on the ELMo embeddings (DeepSeqVec in Fig. 1 and Table 1). However, all models introduced in this work were deliberately designed to be relatively simple to demonstrate the predictive power of *SeqVec*. More sophisticated architectures using *SeqVec* are likely to outperform the approaches introduced here.

When we combined ELMo with evolutionary information for the per-residue predictions, the resulting tool still did not quite achieve top performance (Q3(NetSurfP-2.0)=85.3% vs. Q3(DeepProf + SeqVec)=82.4%, Table 1). This might suggest some limit for the usefulness of ELMo/SeqVec. However, it might also point to the more advanced solutions realized by NetSurfP-2.0 which applies two LSTMs of similar complexity as our entire system (including ELMo) on top of their last step leading to 35M (35 million) free parameters compared to about 244K for DeepProf + SeqVec. Twenty times more free parameters might explain some fraction of the success, however due to limited GPU resources, we could not test how much.

Why did the ELMo-based approach improve more (relative to competition) for per-protein than for per-residue predictions? Per-protein data sets were over two orders of magnitude smaller than those for per-residue (simply because every protein constitutes one sample in the first and protein length samples for the second). It is likely that ELMo helped more for the smaller data sets because the unlabeled data is pre-processed so meaningful that less information needs to be learned by the ANN during per-protein prediction. This view was strongly supported by the t-SNE [52] results (Fig. 2): ELMo apparently had learned enough to realize a very rough clustering into localization and membrane/not.

We picked four particular tasks as proof-of-principle for our ELMo/SeqVec approach. These tasks were picked because recently developed methods implemented deep learning and the associated data sets for training and testing were made publicly available. We cannot imagine why our findings should not hold for other tasks of protein prediction and welcome the community to test our *SeqVec* for their particular tasks. We assume that our findings will be more relevant for small data sets than for large ones. For instance, we assume predictions of inter-residue contacts to improve less, and those for protein binding sites possibly more.

### Good and fast predictions without using evolutionary information

SeqVec predicted secondary structure and protein disorder over 100-times faster on a single 8GB GPU than the top-of-the-line prediction method NetSurfP-2.0 which requires to retrieve evolutionary information summarized in alignments. For some applications, the speedup might outweigh the reduction in performance. Therefore, embedding-based approaches such as *SeqVec* suggested a promising solution toward solving one of the biggest challenges for computational biology: How to efficiently handle the exponentially increasing number of sequences? Here, we showed that relevant information from large unannotated biological databases can be compressed into embeddings that condense and abstract the underlying biophysical principles. These embeddings, essentially the weights of a neural network, help as input to many problems for which smaller sets of annotated data are available (secondary structure, disorder, localization). Although the compression step needed to build the *SeqVec* model is very GPU-intensive, it can be performed in a centralized way using large clusters. After training, the model can be used as input by any consumer hardware. Such solutions are ideal to support researches without access to expensive cluster infrastructure.

### Modeling the language of life?

Our ELMo implementation learned to model a probability distribution over a protein sequence. The sum over this probability distribution constituted a very informative input vector for any machine learning task. It also picked up context-dependent protein motifs without explicitly explaining what these motifs are relevant for. In contrast, tools such as ProtVec will always create the same vectors for a k-mer, regardless of the residues surrounding this k-mer in a particular protein sequence.

Our hypothesis had been that the ELMo embeddings learned from large databases of protein sequences (without annotations) could extract a *probabilistic model of the language of life* in the sense that the resulting system will extract aspects relevant both for per-residue and per-protein prediction tasks. All the results presented here have added independent evidence in full support of this hypothesis. For instance, the three state per-residue accuracy for secondary structure prediction improved by over eight percentage points through ELMo (Table 1: e.g. for CB513: Q3(DeepSeqVec)=76.9% vs. Q3(DeepBLOSUM65)=67.4%, i.e. 9.4 percentage points corresponding to 14% of 67.4), the per-residue MCC for protein disorder prediction also rose (Table 1: e.g. TS115: MCC(DeepSeqVec)=0.591 vs. MCC(DeepBLOSUM65)=0.488, corresponding to 18% of 0.488). On the per-protein level, the improvement over the previously popular tool extracting “meaning” from proteins, ProtVec, was even more substantial (Table 1: localization: Q10(DeepSeqVec-Loc)=68% vs. Q10(DeepProtVec-Loc)=42%, i.e. 62% rise over 42; membrane: Q2(DeepSeqVec-Loc)=86.8% vs. Q2 (DeepProtVec-Loc)=77.6%, i.e. 19% rise over 77.6). We could demonstrate this reality even more directly using the t-SNE [52] results (Fig. 2 and Fig. 3): different levels of complexity ranging from single amino acids, over some localizations, structural features, functions and the classification of membrane/non-membrane had been implicitly learned by SeqVec without any training on such data. Clearly, our ELMo-driven implementation succeeded to model some aspects of the language of life as proxied for proteins.

## Conclusion

We have shown that it is possible to capture and transfer knowledge, e.g. biochemical or biophysical properties, from a large unlabeled data set of protein sequences to smaller, labelled data sets. In this first proof-of-principle, our comparably simple models have already reached promising performance for a variety of per-residue and per-protein prediction tasks obtainable from only single protein sequences as input, that is: without any direct evolutionary information, i.e. without alignments. This reduces the dependence on the time-consuming and computationally intensive calculation of protein profiles, allowing the prediction of per-residue and per-protein features of a whole proteome within less than an hour. For instance, on a single GeForce GTX 1080, the creation of embeddings and predictions of secondary structure and subcellular localization for the whole human proteome took about 32 minutes. Building more sophisticated architectures on top of the proposed SeqVec will increase sequence-based performance further.

Our new SeqVec embeddings may constitute an ideal starting point for many different applications in particular when labelled data are limited. The embeddings combined with evolutionary information might even improve over the best available methods, i.e. enable highquality predictions. Alternatively, they might ease high-throughput predictions of whole proteomes when used as the only input feature. Alignment-free predictions bring speed and improvements for proteins for which alignments are not readily available or limited, such as for intrinsically disordered proteins, for the Dark Proteome, or for particular unique inventions of evolution. The trick was to tap into the potential of Deep Learning through transfer learning from large repositories of unlabeled data by modeling the language of life.

## Methods

### Data

#### UniRef50 training of SeqVec

We trained ELMo on UniRef50 [32], a sequence redundancy-reduced subset of the UniProt database clustered at 50% pairwise sequence identity (PIDE). It contained 25 different letters (20 standard and 2 rare amino acids (U and O) plus 3 special cases describing either ambiguous (B, Z) or unknown amino acids (X); Table SOM_1) from 33M proteins with 9,577,889,953 residues. Each protein was treated as a sentence and each amino acid was interpreted as a single word. We referred to the resulting embedding as to *SeqVec* (*Seq*uence-to-*Vec*tor).

#### Visualization of projection space

The current release of the “Structural Classification Of Proteins” (SCOPe, [53]) database (2.07) contains 14323 proteins at a redundancy level of 40%. Functions encoded by the Enzyme Commission number (E.C., [55]) were retrieved via the “Structure Integration with Function, Taxonomy and Sequence” (SIFTS) mapping [56]. SIFTS allows among other things a residue-level mapping between UniProt and PDB entries and a mapping from PDB identifiers to E.C.s. If no function annotation was available for a protein or if the same PDB identifier was assigned to multiple E.C.s, it was removed from Fig. 3c. Taxonomic identifiers were used to map proteins to one of the 3 kingdoms of life or to viruses. Again, proteins were removed if no such information was available. The number of iterations for the t-SNE projections was set again to 3000 and the perplexity was adjusted (perplexity=5 for Fig. 3a and perplexity=30 for Fig. 3b-3d).

#### Per-residue level: secondary structure & intrinsic disorder (NetSurfP-2.0)

To simplify compatibility, we used the data set published with a recent method seemingly achieving the top performance of the day in secondary structure prediction, namely *NetSurfP-2.0* [45]. Performance values for the same data set exist also for other recent methods such as *Spider3* [57], *RaptorX* [58, 59] and *JPred4* [60]. The set contains 10,837 sequence-unique (at 25% PIDE) proteins of experimentally known 3D structures from the PDB [61] with a resolution of 2.5 Å (0.25 nm) or better, collected by the PISCES server [62]. DSSP [44] assigned secondary structure and intrinsically disordered residues are flagged (residues without atomic coordinates, i.e. REMARK-465 in the PDB file). The original seven DSSP states (+ 1 for unknown) were mapped upon three states using the common convention: [G,H,I] → H (helix), [B,E] → E (strand), all others to O (other; often misleadingly referred to as *coil* or *loop*). As the authors of NetSurfP-2.0 did not include the raw protein sequences in their public data set, we used the SIFTS file to obtain the original sequence. Only proteins with identical length in SIFTS and NetSurfP-2.0 were used. This filtering step removed 56 sequences from the training set and three from the test sets (see below: two from CB513, one from CASP12 and none from TS115). We randomly selected 536 (∼5%) proteins for early stopping (*cross-training*), leaving 10,256 proteins for training. All published values referred to the following three test sets (also referred to as validation set): <u>**TS115**</u> [63]: 115 proteins from high-quality structures (<3Å) released after 2015 (and at most 30% PIDE to any protein of known structure in the PDB at the time); <u>**CB513**</u> [64]: 513 non-redundant sequences compiled 20 years ago (511 after SIFTS mapping); <u>**CASP12**</u> [65]: 21 proteins taken from the CASP12 free-modelling targets (20 after SIFTS mapping; all 21 fulfilled a stricter criterion toward non-redundancy than the two other sets; non-redundant with respect to all 3D structures known until May 2018 and all their relatives). Each of these sets covers different aspects of the secondary structure prediction problem: CB513 and TS115 only use structures determined by X-ray crystallography, applying similar cutoffs with respect to redundancy (30%) and resolution (2.5-3.0Å). While they serve as a good proxy for a baseline performance, CASP12 might better reflect the true generalization capability for unseen proteins including NMR and Cryo-EM as means of structure determination. Also, the strict redundancy reduction based on publication date reduces the bias towards well studied families. Nevertheless, toward our objective of establishing a proof-of-principle, these sets sufficed. All test sets had fewer than 25% PIDE to any protein used for training and cross-training (ascertained by the *NetSurfP-2.0* authors). To compare methods using evolutionary information and those using our new word embeddings, we took the *HHblits* profiles published along with the NetSurfP-2.0 data set.

#### Per-protein level: localization & membrane proteins (DeepLoc)

Localization prediction was trained and evaluated using the *DeepLoc* data set [46] for which performance was measured for several methods, namely: LocTree2 [66], MultiLoc2 [67], SherLoc2 [68], CELLO [69], iLoc-Euk [51], WoLF PSORT [70] and YLoc [71]. The data set contained proteins from UniProtKB/Swiss-Prot [48] (release: 2016_04) with experimental annotation (code: ECO:0000269). The *DeepLoc* authors mapped these to ten classes, removing all proteins with multiple annotations. All these proteins were also classified into *water-soluble* or *membrane-bound* (or as *unknown* if the annotation was ambiguous). The resulting 13,858 proteins were clustered through PSI-CD-HIT [72, 73] (version 4.0; at 30% PIDE or Eval<10^−6^). Adding the requirement that the alignment had to cover 80% of the shorter protein, yielded 8,464 clusters. This set was split into training and testing by using the same proteins for testing as the authors of DeepLoc. The training set was randomly sub-divided into 90% for training and 10% for determining early stopping (cross-training set).

### ELMo terminology

<u>One-hot encoding</u> (also known as *sparse encoding*) assigns each word (referred to as <u>token</u> in NLP) in the vocabulary an integer N used as the Nth component of a vector with the dimension of the vocabulary size (number of different words). Each component is binary, i.e. either 0 if the word is not present in a sentence/text or 1 if it is. This encoding drove the first application of machine learning that clearly improved over all other methods in protein prediction [1-3]. <u>TF-IDF</u> represents tokens with the product of “frequency of token in data set” times “inverse frequency of token in document”. Thereby, rare tokens become more relevant than common words such as “the” (so called *stop words*). This concept resembles that of using k-mers for database searches [33], clustering [74], motifs [75, 76], and prediction methods [66, 70, 77-81]. <u>Context-insensitive word embedding</u> replaced expert features, such as TF-IDF, by algorithms that extracted such knowledge from unlabeled corpus such as Wikipedia, by either predicting the neighboring words, given the center word (skip-gram) or vice versa (CBOW). This became known in *Word2Vec* [49] and showcased for computational biology through *ProtVec* [49, 82]. More specialized implementations are *mut2vec* [83] learning mutations in cancer, and *phoscontext2vec* [84] identifying phosphorylation sites. The performance of context-insensitive approaches was pushed to its limits by adding sub-word information (FastText [85]) or global statistics on word co-occurance (GloVe [86]). <u>Context-sensitive word embedding</u> started a new wave of word embedding techniques for NLP in 2018: the particular embedding renders the meaning of the phrase “*paper tiger”* dependent upon the context. Popular examples like ELMo [41] and Bert [87] have achieved state-of-the-art results in several NLP tasks. Both require substantial GPU computing power and time to be trained from scratch. However, in this work we focused on ELMo as it allows processing of sequences of variable length. This ELMo model consists of a single CharCNN [88] over the characters in a word and two layers of bidirectional LSTMs that introduce the context information of surrounding words. The CharCNN transforms all characters within a single word via an embedding layer into vector space and runs multiple CNNs of varying window size (here: ranging from 1 to 7) and number of filters (here: 32, 64, …, 1024). In order to obtain a fixed-dimensional vector for each word, regardless of its length, the output of the CNNs is max-pooled and concatenated. The first bi-directional LSTM operates directly on the output of the CharCNN, while the second LSTM layer takes the output of the first LSTM as input. As described in the original ELMo publication, the weights of the forward and the backward model are shared in order to reduce the memory overhead of the model and to combat overfitting. Even though, the risk of overfitting is small due to the high imbalance between number of trainable parameters (93M) versus number of tokens (9.3B), dropout at a rate of 10% was used to reduce the risk of overfitting. This model is trained to predict the next word given all previous words in a sentence. To the best of our knowledge, the context-sensitive ELMo has not been adapted to protein sequences, yet.

### ELMo adaptation

In order to allow more flexible models and easily integrate into existing solutions, we have used and generated ELMo as word embedding layers. No fine-tuning was performed on task-specific sequence sets. Thus, researchers could just replace their current embedding layer with our model to boost their task-specific performance. Furthermore, it will simplify the development of custom models that fit other use-cases. The embedding model takes a protein sequence of arbitrary length and returns 3076 features for each residue in the sequence. These 3076 features were derived by concatenating the outputs of the three internal layers of ELMo (1 CharCNN-layer, 2 LSTM-layers), each describing a token with a vector of length 1024. For simplicity, we summed the components of the three 1024-dimensional vectors to form a single 1024-dimensional feature vector describing each residue in a protein. In order to demonstrate the general applicability of *SeqVec*, we neither fine-tuned the model on a specific prediction task, nor optimized the combination of the three internal layers. Instead, we used the standard ELMo configuration [41] with the following changes: (i) reduction to 28 tokens (20 standard and 2 rare (U,O) amino acids + 3 special tokens describing ambiguous (B,Z) or unknown (X) amino acids + 3 special tokens for ELMo indicating padded elements (‘<MASK>’) or the beginning (‘<S>’) or the end of a sequence (‘</S>’)), (ii) increase number of unroll steps to 100, (iii) decrease number of negative samples to 20, (iv) increase token number to 9,577,889,953.

### SeqVec 2 prediction

<u>On the per-residue level</u>, the predictive power of the new *SeqVec* embeddings was demonstrated by training a small two-layer Convolutional Neural Network (CNN) in PyTorch using a specific implementation [89] of the ADAM optimizer [90], cross-entropy loss, a learning rate of 0.001 and a batch size of 128 proteins. The first layer (in analogy to the sequence-to-structure network of earlier solutions [1, 2]) consisted of 32-filters each with a sliding window-size of w=7. The second layer (structure-to-structure [1, 2]) created the final predictions by applying again a CNN (w=7) over the output of the first layer. These two layers were connected through a rectified linear unit (ReLU) and a dropout layer [91] with a dropout-rate of 25% (Fig. 4, left panel). This simple architecture was trained independently on six different types of input, each with a different number of free parameters. (i) <u>DeepProf (14,000=14k free parameters)</u>: Each residue was described by a vector of size 50 which included a one-hot encoding (20 features), the profiles of evolutionary information (20 features) from HHblits as published previously [45], the state transition probabilities of the Hidden-Markov-Model (7 features) and 3 features describing the local alignment diversity. (ii) Deep<u>SeqVec (232k free parameters):</u> Each protein sequence is represented by the output from SeqVec. The resulting embedding described each residue as a 1024-dimensional vector. <u>(iii) DeepProf+SeqVec (244k free parameters):</u> This model simply concatenated the input vectors used in (i) and (ii). <u>(iv) DeepProtVec (25k free parameters):</u> Each sequence was split into overlapping 3-mers each represented by a 100-dimensional ProtVec [47]. <u>(v) DeepOneHot (7k free parameters):</u> The 20 amino acids were encoded as one-hot vectors as described above. Rare amino acids were mapped to vectors with all components set to 0. Consequently, each protein residue was encoded as a 20-dimensional one-hot vector. <u>(vi) DeepBLOSUM65 (8k free parameters):</u> Each protein residue was encoded by its BLOSUM65 substitution matrix [92]. In addition to the 20 standard amino acids, BLOSUM65 also contains substitution scores for the special cases B, Z (ambiguous) and X (unknown), resulting in a feature vector of length 23 for each residue.

**Figure 4:**
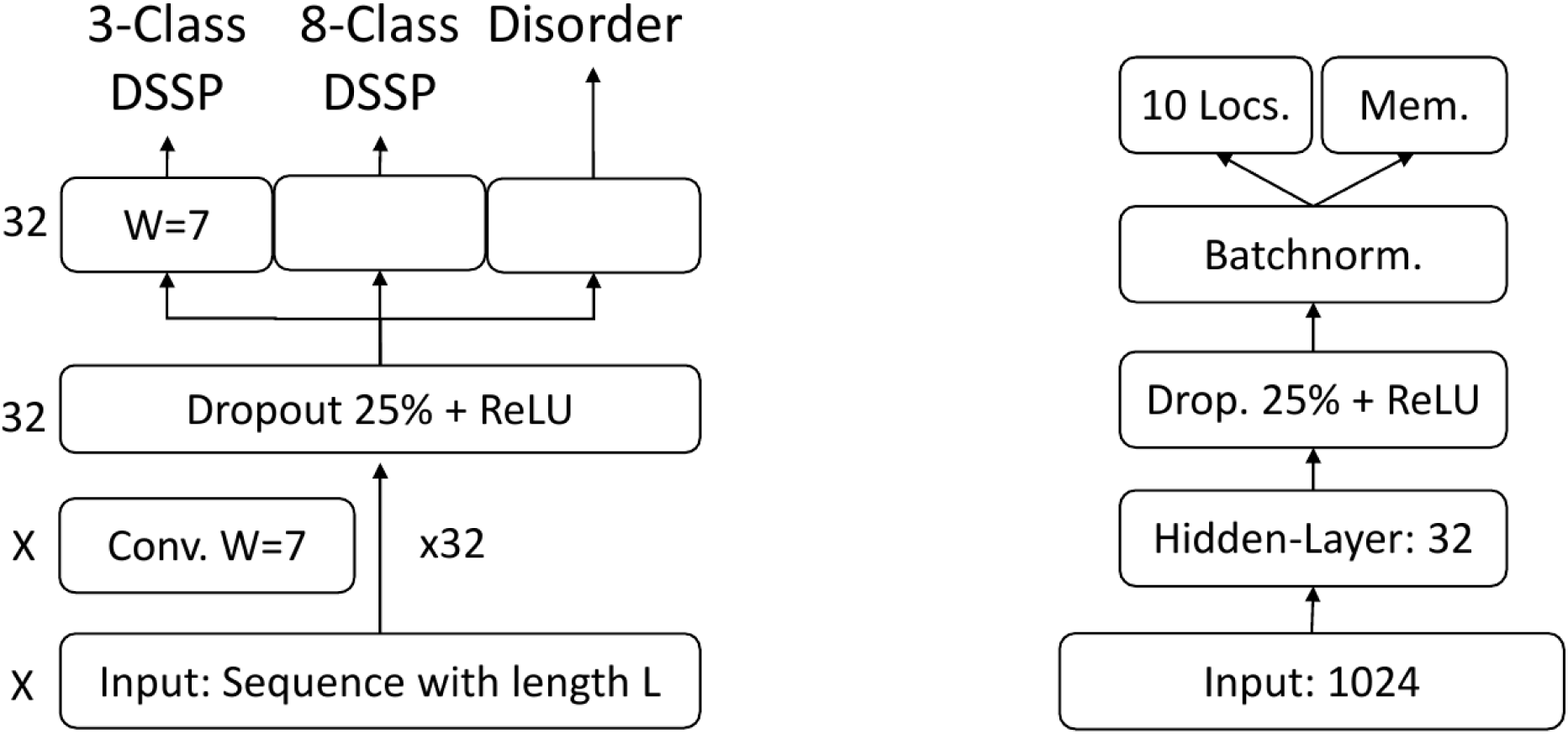
On the left the architecture of the model used for the per-residue level predictions (secondary structure and disorder) is sketched, on the right that used for per-protein level predictions (localization and membrane/not membrane). The ‘X’, on the left, indicates that different input features corresponded to a difference in the number of input channels, e.g. 1024 for *SeqVec* or 50 for profile-based input. The letter ‘W’ refers to the window size of the corresponding convolutional layer (W=7 implies a convolution of size 7×1).

<u>On the per-protein level</u>, a simple feed-forward neural network was used to demonstrate the power of the new embeddings. In order to ensure equal-sized input vectors for all proteins, we averaged over the embeddings of all residues in a given protein resulting in a 1024-dimensional vector representing any protein in the data set. ProtVec representations were derived the same way, resulting in a 100-dimensional vector. These vectors (either 100- or 1024 dimensional) were first compressed to 32 features, then dropout with a dropout rate of 25%, batch normalization [93] and a rectified linear Unit (ReLU) were applied before the final prediction (Fig. 4, right panel). In the following, we refer to the models trained on the two different input types as (i) <u>DeepSeqVec-Loc (33k free parameters)</u>: average over SeqVec embedding of a protein as described above and (ii) <u>DeepProtVec-Loc (320 free parameters)</u>: average over ProtVec embedding of a protein. We used the following hyper-parameters: learning rate: 0.001, Adam optimizer with cross-entropy loss, batch size: 64. The losses of the individual tasks were summed before backpropagation. Due to the relatively small number of free parameters in our models, the training of all networks completed on a single Nvidia GeForce GTX1080 within a few minutes (11 seconds for DeepSeqVec-Loc, 15 minutes for DeepSeqVec).

### Evaluation measures

To simplify comparisons, we ported the evaluation measures from the publications we derived our data sets from, i.e. those used to develop *NetSurfP-2.0* [45] and *DeepLoc* [46]. All numbers reported constituted averages over all proteins in the final test sets. This work aimed at a proof-of-principle that the *SeqVec* embedding contain predictive information. In the absence of any claim for state-of-the-art performance, we did not calculate any significance values for the reported values.

#### Per-residue performance

Toward this end, we used the standard three-state per-residue accuracy (Q3=percentage correctly predicted in either helix, strand, other [1]) along with its eight-state analog (Q8). Predictions of intrinsic disorder were evaluated through the Matthew’s correlation coefficient (MCC [94]) and the False-Positive Rate (FPR) representative for tasks with high class imbalance. For completeness, we also provided the entire confusion matrices for both secondary structure prediction problems (Fig. SOM_2). Standard errors were calculated over the distribution of each performance measure for all proteins.

#### Per-protein performance

The predictions whether a protein was membrane-bound or water-soluble were evaluated by calculating the two-state per set accuracy (Q2: percentage of proteins correctly predicted), and the MCC. A generalized MCC using the Gorodkin measure [95] for K (=10) categories as well as accuracy (Q10), was used to evaluate localization predictions. Standard errors were calculated using 1000 bootstrap samples, each chosen randomly by selecting a sub-set of the predicted test set that had the same size (draw with replacement).

## Abbreviations

1D: one-dimensional – information representable in a string such as secondary structure or solvent accessibility;
3D: three-dimensional;
3D structure: three-dimensional coordinates of protein structure;
MCC: Matthews-Correlation-Coefficient;
RSA: relative solvent accessibility;

## Availability

The pre-trained ELMo-like SeqVec model and a description on how to implement the embeddings into existing methods can be found here: https://github.com/Rostlab/SeqVec. Accessed 10^th^ September 2019.

Predictions on secondary structure, disorder and subcellular localization based on SeqVec can be accessed under: https://embed.protein.properties. Accessed 10^th^ September 2019.

The NetSurfP-2.0 data set [45] used for the evaluation of SeqVec on the task of secondary structure and disorder prediction are publicly available under: http://www.cbs.dtu.dk/services/NetSurfP/. Accessed 10^th^ September 2019.

The DeepLoc data set [46] used for the evaluation of SeqVec on the task of subcellular localization prediction are publicly available under: http://www.cbs.dtu.dk/services/DeepLoc/data.php. Accessed 10^th^ September 2019.

## Authors contributions

AE and MH suggested to use ELMo for modeling protein sequences. AE adopted and trained ELMo. MH evaluated SeqVec embeddings on different data sets and tasks. YW helped with discussions about natural language processing. CD implemented the web-interface which allows to access and visualize the predictions. DN helped with various problems regarding the code. FM and BR helped with the design of the experiment and to critically improve the code. FM and BR helped with the design of the experiment and to critically improve the manuscript. MH and AE drafted the manuscript and the other authors provided feedback. All authors read and approved the final manuscript.

## Acknowledgements

The authors thank primarily Tim Karl for invaluable help with hardware and software and Inga Weise for support with many other aspects of this work. Last, not least, thanks to all those who deposit their experimental data in public databases, and to those who maintain these databases.

