## Supplementary Online Material for "Modeling the language of life – Deep Learning Protein Sequences"

### Supporting online material for: Modeling the Language of Life – Deep Learning Protein Sequences

#### Table of Contents for Supporting Online Material

#### Short description of Supporting Online Material

The evolution of the model's uncertainty (or perplexity) when predicting the next token during training is shown in Fig. SOM_1. The vocabulary and the number of occurrences of the tokens which are used to train ELMo are shown in Table SOM_1.

Confusion matrices for predictions on the level of residues (Figure SOM_2) and on the level of whole proteins (Figure SOM_3) are given in the following.

### SOM: Modelling the language of life

#### Fig. SOM_1: ELMo perplexity

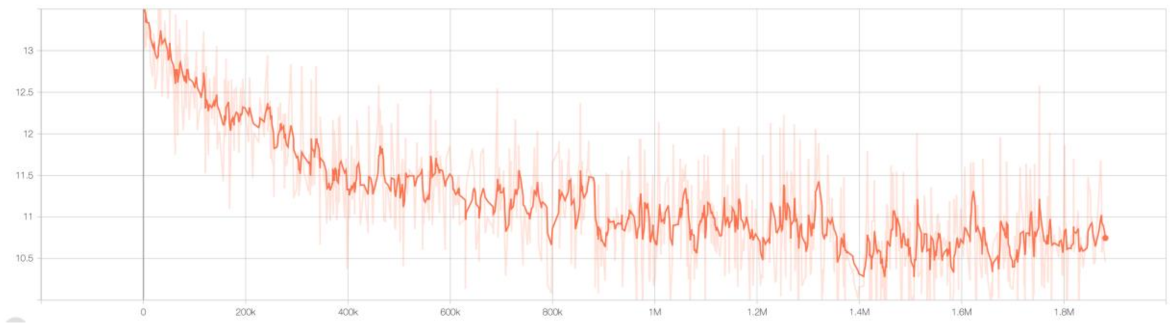

**Fig. SOM_1: ELMo perplexity.** The perplexity defines the uncertainty of a model when predicting the next token (here: amino acid), given all previous tokens in a sequence. Lower values indicate less uncertainty. This measure can be used to monitor the training progress (y-axis: perplexity) over time (x-axis: number of training steps). Here, the learning progress of the proposed ELMo-based SeqVec is shown while being trained on UniRef50.

### Fig. SOM_2: Confusion matrices for per-protein predictions using SeqVec

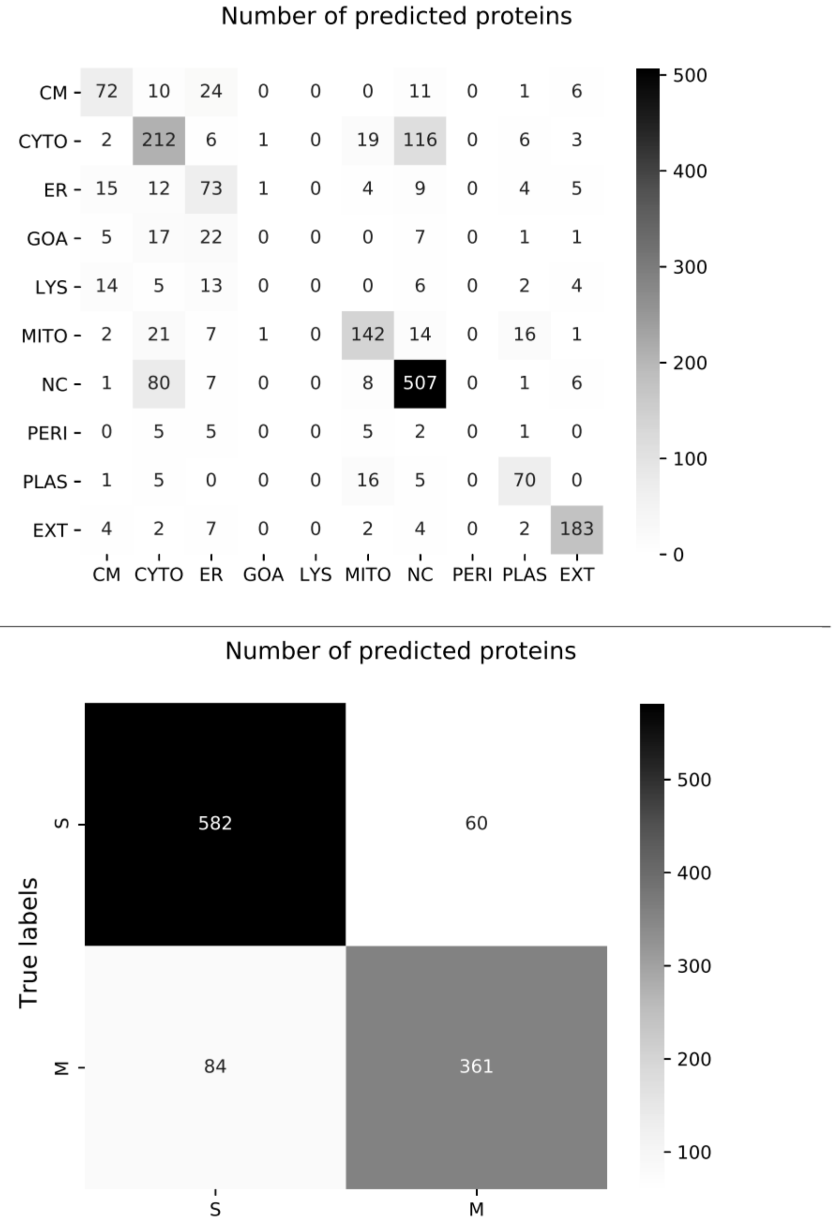

**Fig. SOM_2: Confusion matrices for per-protein predictions using only SeqVec.** The confusion matrix for localization prediction is shown in the upper row, while results for membrane-bound versus water-soluble are given in the lower row. Again, each confusion matrix summarizes true labels (rows) and predictions (columns). Localizations are abbreviated for simplicity (CM=cell membrane, CYTO=cytoplasm, ER=endoplasmic reticulum, GOA=golgi apparatus, LYS=Lysosome/Vacuole, MITO=mitochondrion, NC=nucleus, PERI=peroxisome, PLAS=plastid, EXT=extracellular).

#### Fig. SOM_3: Confusion matrices for secondary structure predictions of DeepSeqVec

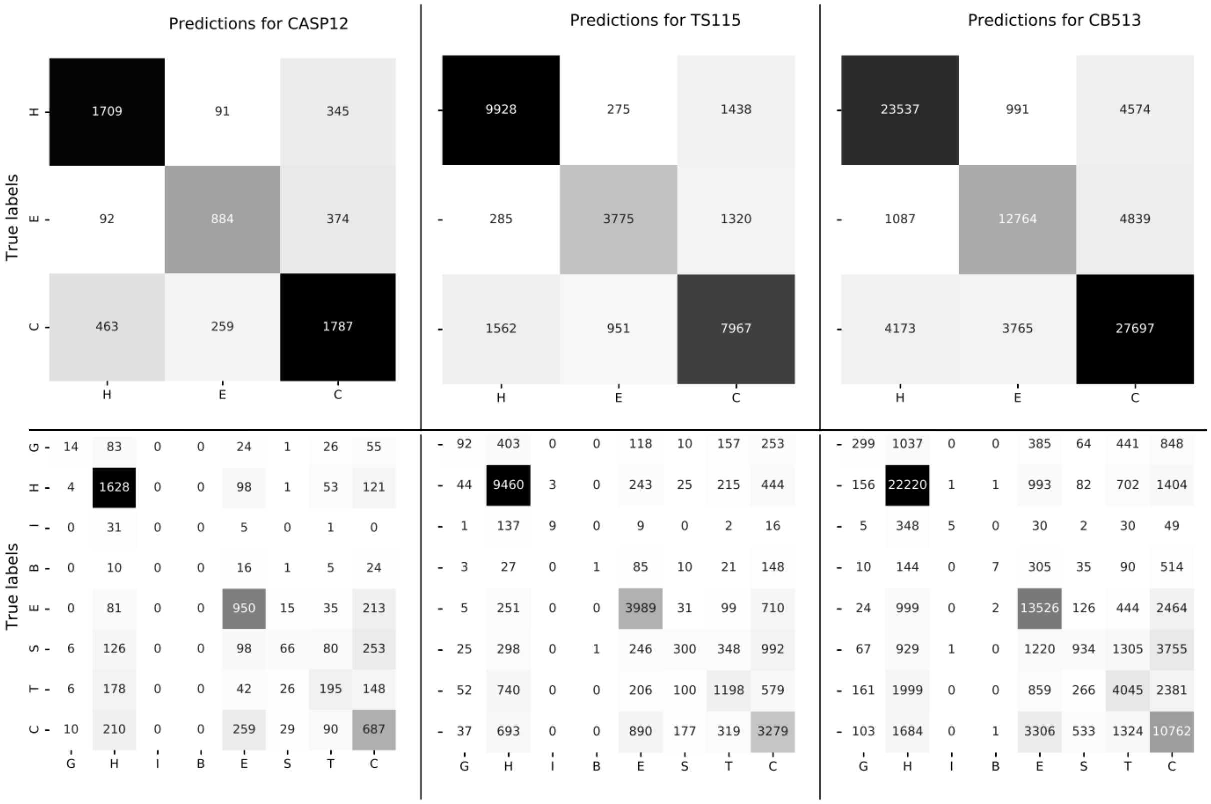

**Fig. SOM_3: Confusion matrices for secondary structure predictions of DeepSeqVec.** The confusion matrices for 3-state secondary structure prediction are given in the upper row, 8-state confusion matrices in the lower row. The columns refer to the three different test sets: CASP12, TS115, CB513. Each matrix summarizes the true labels (rows) and the predictions (columns) for each of the sets or tasks. Numbers on the diagonal reflect correct predictions.

**Table SOM_1: Amino acid occurrences in UniRef50 ◊**

| Amino acid (single letter) | Number of occurrences | % of data | Comment |
|---|---|---|---|
| L | 918255239 | 9.6 | Leucine |
| A | 815091587 | 8.5 | Alanine |
| S | 721399187 | 7.5 | Serine |
| G | 651415980 | 6.8 | Glycine |
| V | 620159476 | 6.5 | Valine |
| E | 590517634 | 6.2 | Glutamic Acid |
| R | 557775181 | 5.8 | Arginine |
| T | 549632104 | 5.7 | Threonine |
| I | 530954378 | 5.5 | Isoleucine |
| D | 525972161 | 5.5 | Aspartic Acid |
| K | 494682943 | 5.2 | Lysine |
| P | 479898431 | 5.0 | Proline |
| N | 405230829 | 4.2 | Asparagine |
| F | 374988414 | 3.9 | Phenylalanine |
| Q | 374551957 | 3.9 | Glutamine |
| Y | 284178589 | 3.0 | Tyrosine |
| H | 210995340 | 2.2 | Histidine |
| M | 207346116 | 2.2 | Methionine |
| C | 135532698 | 1.4 | Cysteine |
| W | 123527183 | 1.3 | Tryptophan |
| X | 5778293 | 0.06 | Any amino acid |
| B | 3780 | 4e-5 | D or N |
| Z | 1363 | 1e-5 | E or Q |
| U | 1043 | 1e-5 | Selenocysteine |
| O | 47 | 5e-7 | Pyrrolysine |
| Total | 9577889953 | 100 | |

◊ Given are the occurrences of all 20 common plus 2 rare amino acids (U and O) and 3 symbols for special cases (B means either D or N, Z means either E or Q and X means that the residue is unknown) which were used to train ELMo on all 33M proteins from UniRef50 (sorted in decreasing order).
